# Photoperiodicity in Glucose Metabolism in the Human Brain

**DOI:** 10.1101/2024.08.26.609495

**Authors:** Kyoungjune Pak, Seunghyeon Shin, Keunyoung Kim, Jihyun Kim, Hyun-Yeol Nam, Lauri Nummenmaa, Pirjo Nuutila, Xingdang Liu, Lihua Sun

## Abstract

Photoperiodicity in the human brain function, which is a critical factor for social well-being, has been widely debated. In this study, 432 healthy males underwent fasting-state brain [^18^F]fluorodeoxyglucose positron emission tomography (PET) scanning twice: first at the baseline and then at the 5-year follow-up. We analyzed the effect of day length on brain glucose uptake separately for the baseline and follow-up studies and examined changes in glucose consumption as a function of the day length deviation for each participant between the repeated PET scans. Glucose uptake in the cuneus was consistently predicted by the day length on the day of scanning and by within-participant day length deviations. This longitudinal large-scale PET study provides a landmark evidence for photoperiodicity in glucose metabolism in the human brain. The cuneus may be an essential part of the visual cortex, translating environmental photoperiodic changes into temporal cues that influence cognitive function and social behavior.

**Significance statement:** Photoperiodicity in the human brain function has been widely debated. The current study provides a landmark evidence in this regard by demonstrating how the photoperiod shapes glucose metabolism in the brain of healthy males, highlighting the crucial role of the cuneus in processing visual information related to photoperiodic changes in the environment. Understanding photoperiodicity in the function of the human brain offers insights into how humans adapt to their environments for social well-being and underscores the potential health implications of changes in the exposure of natural light.

## Introduction

The historical debate on photoperiodicity in the function and physiology of the human brain has evolved, as evidenced by the increasing modern neuroscientific support for its existence (1-6). The human visual pathway begins primarily in the retina, where rod and cone cells detect and convert light into electrical signals. These signals then travel through the optic tract to the lateral geniculate nucleus in the thalamus, which relays them to the primary visual cortex (V1) for further processing. Modern neuroscience has significantly advanced our understanding of visual information processing in the V1, including the roles of the dorsal and ventral pathways in movement and object recognition (7, 8). However, how the visual cortex may encode information regarding the duration of light (i.e., photoperiod) and translate this into temporal cues that guide seasonal rhythms of brain functions remains unclear.

Researchers supporting the concept of photoperiodicity emphasize the evolutionary significance of biological clocks and fundamental role of natural light in regulating various physiological processes. Throughout human history, exposure to the natural light-dark cycle has been a consistent environmental cue that guides circadian rhythms. Proponents of photoperiodicity further contend that disruptions to this natural cycle, particularly through increased exposure to artificial light at inappropriate times, can lead to a range of health issues, such as sleep disturbances, mood disorders, and disruptions in hormonal regulation (1-3, 9). Accordingly, understanding and respecting the inherent connection between brain physiology and the natural light-dark cycle is essential for promoting optimal health and well-being (10).

To this end, we explored the potential photoperiodicity in humans based on a unique database in which 432 healthy males underwent longitudinal, repeated [^18^F]fluorodeoxyglucose (FDG) positron emission tomography (PET) measurements of brain glucose consumption, along with blood sample features registered on the same days as the PET scans. Glucose serves as the main energy source for the brain and body, and dysregulation of brain glucose consumption is involved in almost all mental health disorders. Fluctuations in the supply of glucose to the brain and body affect cognition and behavior (11, 12). Abnormal glucose uptake has been frequently linked to psychiatric disorders, such as depression (13), autism, and schizophrenia (14). Additionally, diabetes, which is associated with atypical brain glucose metabolism, is known to increase the risk of depression, anxiety, and eating disorders (15). Therefore, a deep knowledge of the biological rhythms of brain glucose consumption is essential for social well-being and mental health.

In the present study, the day length on the day of the brain PET scan was first used as a predictor of brain glucose uptake (GU) for both the baseline and 5-year follow-up studies. Furthermore, within-participant changes in brain glucose consumption were investigated as a function of day length deviation between repeated brain PET scans. We believe that these data are sufficiently powered to provide evidence that support or refute the presence of photoperiodicity in the human brain.

## Methods

### Subjects

We retrospectively analyzed the data of 473 healthy males who underwent a health checkup program at the Samsung Changwon Hospital Health Promotion Center in 2013 (baseline) and 2018 (5-year follow-up). After excluding those with neuropsychiatric disorders (n=8) or malignancies (n=3) and those with missing anthropometric and body composition measurements (n=30), 432 healthy males were included in both the baseline (mean 42.8, range 38–50 years) and follow-up (mean 47.9, range 43–55 years) studies. The distribution of the day length data is shown in **Fig. 1**.

**Figure 1.**
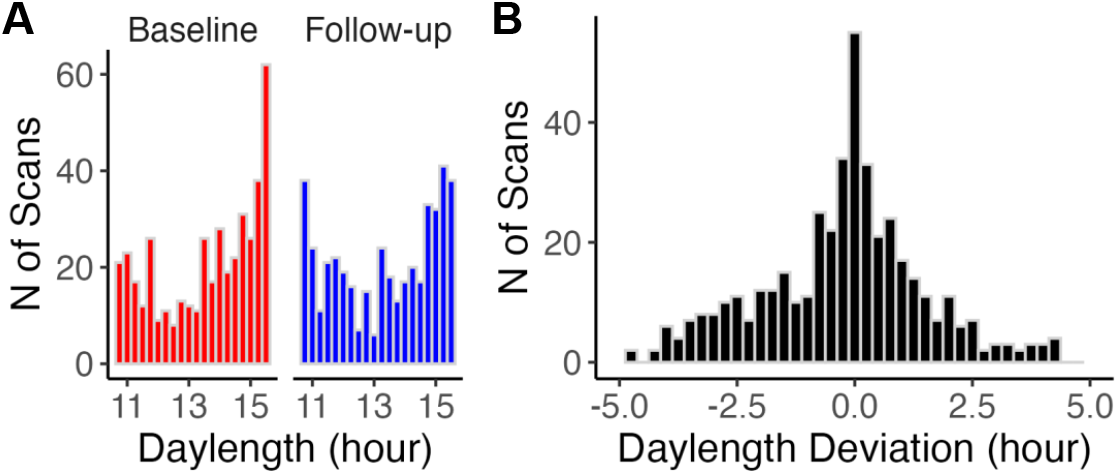
Distribution of Day length. **A)** Distribution of day length in the baseline and follow-up studies. **B)** Distribution of within-participant deviation in day length between repeated scans. One bin represents a day length of 0.25 hours. The distributions of other relevant variables are shown in **Supplementary Figure S1**. Further details on the participants are provided in **Supplementary Tables S1 and S2**.

Participants of this study were included in a previous study that investigated the effects of aging on brain glucose metabolism (16). The health checkup program included 1) a brain [^18^F]FDG PET scan, 2) blood sampling, and 3) administration of depression and stress questionnaires on the same day. In addition, sociodemographic information, including marital status and educational level, was recorded. The study protocol was approved by the Institutional Review Board of Changwon Samsung Hospital. The requirement for informed consent was waived because of the retrospective nature of the study.

### Brain [^18^F]FDG PET acquisition and Image analysis

The participants were asked to avoid strenuous exercise for 24 hours and fast for at least 6 hours before the PET study. PET/computed tomography (CT) was performed 60 minutes after the injection of [^18^F]FDG (3.7 MBq/kg) using a Discovery 710 PET/CT scanner (GE Healthcare, Waukesha, WI, USA). Continuous spiral CT was performed with a tube voltage of 120 kVp and tube current of 30–180 mAs. PET scans were obtained in a three-dimensional mode with a full width at half maximum of 5.6 mm and reconstructed using an ordered-subset expectation maximization algorithm. PET scans were spatially normalized to the MNI space using PET templates from SPM5 (University College of London, UK) with pmod version 3.6 (PMOD Technologies LLC, Zurich, Switzerland). An Automated Anatomical Labeling 2 (AAL2) atlas (17) was used to define regions of interest (ROIs), including the frontal pole, insula, orbitofrontal cortex, anterior cingulate cortex, posterior cingulate cortex, precuneus, cuneus, amygdala, striatum (caudate and putamen), hippocampus, superior temporal gyrus, middle temporal gyrus, pre- and postcentral cortex, and cerebellum. These 15 ROIs were selected based on their roles in emotional processing and social cognition (18). The mean uptake of each ROI was scaled to the mean global cortical uptake of each individual and was defined as the standardized uptake value ratio (SUVR). For full-volume analysis, the statistical threshold was set at the cluster level and corrected with a false discovery rate (p < 0.05) in a regression model (corrected for age and BMI) after smoothing SUVR images with a Gaussian kernel of full width at half maximum of 8 mm (Statistical Parametric Mapping 12, Wellcome Centre for Human Neuroimaging, UCL, London, UK).

### Anthropometric measurements and blood tests

The height (cm) and weight (kg) of participants were measured, and body mass index (BMI) was calculated as weight divided by the square of height (kg/m^2^). Blood samples were collected from the antecubital veins of each participant. Fasting plasma glucose (mg/dL), insulin (μIU/mL), serum sodium (mmol/L), and serum potassium (mmol/L) levels, as well as red blood cells (RBC), white blood cells (WBC), and platelet counts were measured.

### Depression and Stress Questionnaires

The participants completed the depression and stress questionnaires. The Center for Epidemiologic Studies Depression Scale (CES-D) consists of 20 self-reported items with scores ranging from 0 to 60, for which higher scores indicate a more severe depression (19). The stress questionnaire for the Korean National Health and Nutrition Examination Survey (KNHANES) consists of 9 self-reported items with scores ranging from 9 to 45; higher scores indicate more stress (20).

### Statistical analysis

#### Full-volume voxel-level analysis

For each participant, the day length was calculated as the sum of daytime and civil twilight on the day of the brain PET scan, as previously described (5, 21). Civil twilight comprises morning civil twilight, which begins when the geometric center of the sun is 6° below the horizon and ends at sunrise, and evening civil twilight, which begins at sunset and ends when the geometric center of the sun reaches 6° below the horizon. Calculation was performed using the R package “suncalc;” the calculations were based on the geographic location of Changwon city, Republic of Korea (latitude = 35.2271° N; longitude = 128.6811° E). In the analysis, age and body mass index (BMI) of the participants were used as nuisance covariates. Data from the two different years of study were analyzed separately.

#### Region of interest (ROI) analysis

Data were analyzed using linear effects regression with the R statistical software (version 4.4.1). Data from the two different years were analyzed separately using day length, age, and BMI as regressors. ROI data were log-transformed, and regressors were standardized (z-score transformed), as previously described (21). In addition, within-participant variations in GU were analyzed using the within-participant deviations of day length and change in BMI as regressors. The within-participant variation in the ROI data and independent variables were standardized in the statistical analysis.

#### Analysis of physiological measures

Anthropometric measures (including BMI) from the baseline and follow-up studies were first analyzed separately, using day length and age as regressors. Additionally, within-participant variations in blood tests were analyzed using the within-participant deviations in day length and change in BMI as regressors. Both the dependent and independent variables were standardized in these analyses.

#### Comparison of variables between the baseline and follow-up

The Wilcoxon rank test and chi-squared test were used to compare variables between the baseline and follow-up studies. Statistical analysis was performed using the R Statistical Software (R Foundation for Statistical Computing).

## Results

### Participant characteristics

A total of 432 individuals were included as participants in both the baseline and follow-up studies. During the 5-year follow-up, BMI (p<0.0001), RBC count (p=0.0012), platelet count (p<0.0001), glucose level (p<0.0001), and insulin level (p<0.0001) increased, whereas WBC counts (p<0.0001) and stress scores (p=0.0021) decreased. The characteristics of the participants in the baseline and follow-up studies are summarized in **Supplementary Table S1**.

### Analysis of brain PET scans at baseline and follow-up

In both the baseline and follow-up PET scans, we found consistent effects of day length on GU in brain clusters spanning the superior and middle frontal gyri, precentral area, precuneus, cuneus, middle temporal gyrus, and middle occipital gyrus, where GU levels were positively correlated with day length (**Fig. 2**). In contrast, GU levels were negatively correlated with day length in the parahippocampal area, amygdala, hypothalamus, fusiform gyrus, lower middle frontal gyrus, and cerebellum.

**Figure 2.**
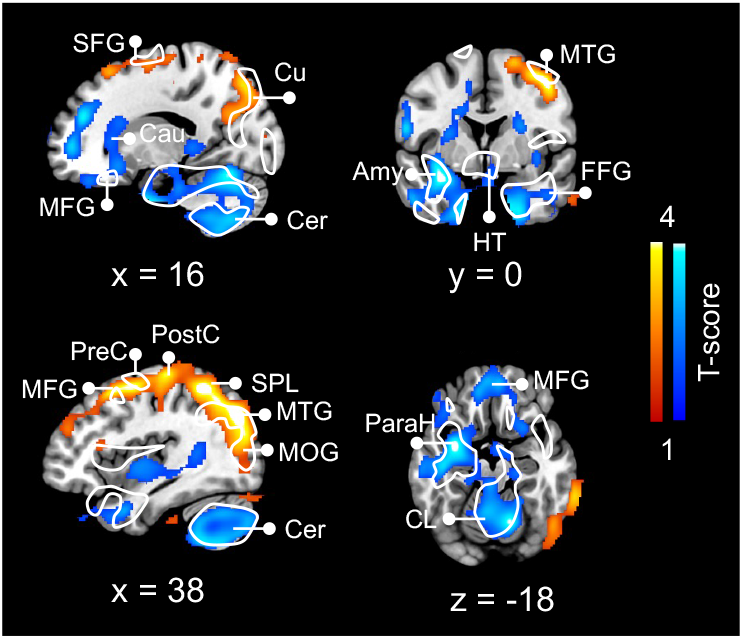
Brain regions where glucose uptake is sensitive to day length. Baseline measures demonstrate positive (hot color) and negative (cold color) relationship with day length, whereas contoured regions show similar effects in the follow-up scans (see **Supplementary Figure S2**). Data are FDR cluster-level corrected at p < 0.05. Amy, amygdala; Cer, cerebellum; CL, cerebellar lingual gyrus; Cu, cuneus; FFG, fusiform gyrus; HT, hypothalamus; MFG, middle frontal gyrus; MOG, middle occipital gyrus; MTG, middle temporal gyrus; ParaH parahippocampus; PostC, postcentral gyrus; PreC, precentral gyrus; SFG, superior frontal gyrus; SPL, superior parietal lobe.

These findings were consistent with the ROI analysis results, showing that in both the baseline and follow-up PET scans, day length influenced regional GU in the cuneus, orbitofrontal cortex, hippocampus, and cerebellum (**Fig. 3**). Specifically, longer day length was associated with increased GU in the cuneus and orbitofrontal cortex and reduced GU in the hippocampus and cerebellum.

**Figure 3.**
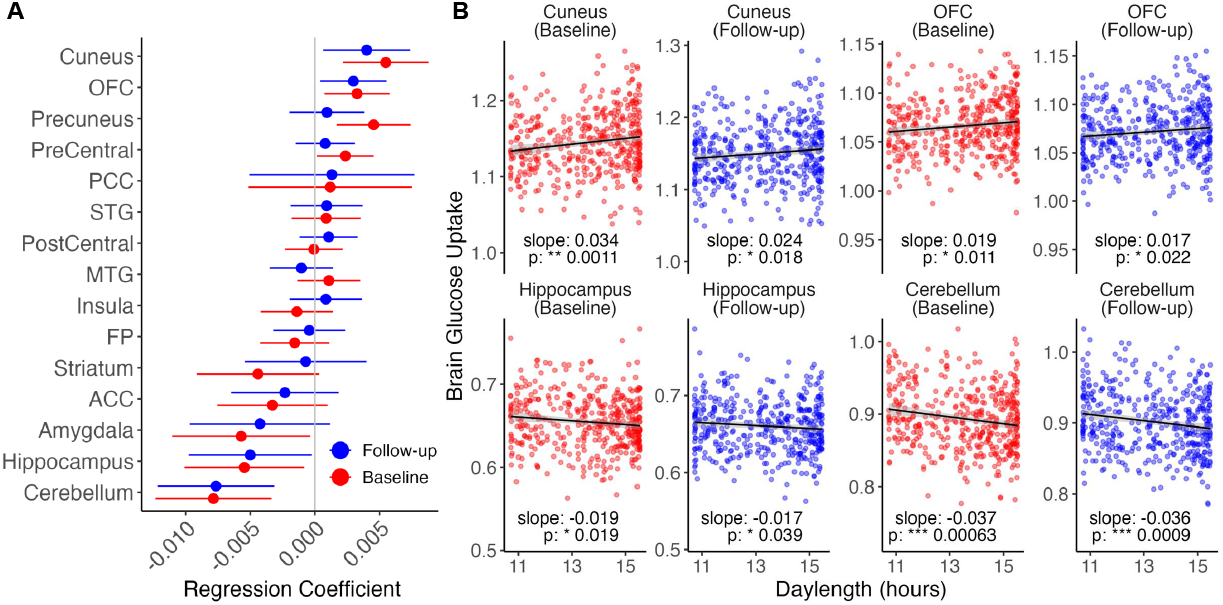
Brain regions where glucose uptake is sensitive to day length in both baseline and follow-up scans. **A)** 95% confidence intervals for individual ROIs where glucose uptake is affected by day length. Log-scale regression coefficients are plotted. **B)** Dot plots of selected ROIs showing the impact of day length on regional glucose uptake. Black line shows the least-squares (LS) regression line (y ~ daylength + BMI + age) predicting glucose uptake, and the shaded area shows 95% confidence intervals. * = p <0.05, ** = p <0.01, *** = p <0.001. ACC, anterior cingulate cortex; BMI, body mass index; FP, frontal pole; OFC, orbitofrontal cortex; PCC, posterior cingulate cortex; ROI, region of interest.

We previously reported that age and BMI are effectors of brain glucose consumption in each analysis (16). Age, BMI, and day length were not correlated in the baseline as well as follow-up studies (**Supplementary Figure S3**).

### Repeated-measures analysis

We further analyzed the within-participant changes in regional GU (i.e., the increase or decrease in the follow-up scan compared with that in the baseline scan) to investigate whether this change was explained by day length deviations between the repeated scans. The day length in the follow-up study was longer than that in the baseline study in 196 participants (45.4%) (**Supplementary Table S2)**. Brain PET scans performed on the day of longer or shorter daylength had comparable age distributions (45.64 ± 4.32 versus 45.08 ± 4.41 years). These findings confirmed that longer day length was associated with increased GU in the cuneus and orbitofrontal cortex (**Fig. 4**). Additionally, increased day length was associated with reduced GU in the anterior cingulate cortex.

**Figure 4.**
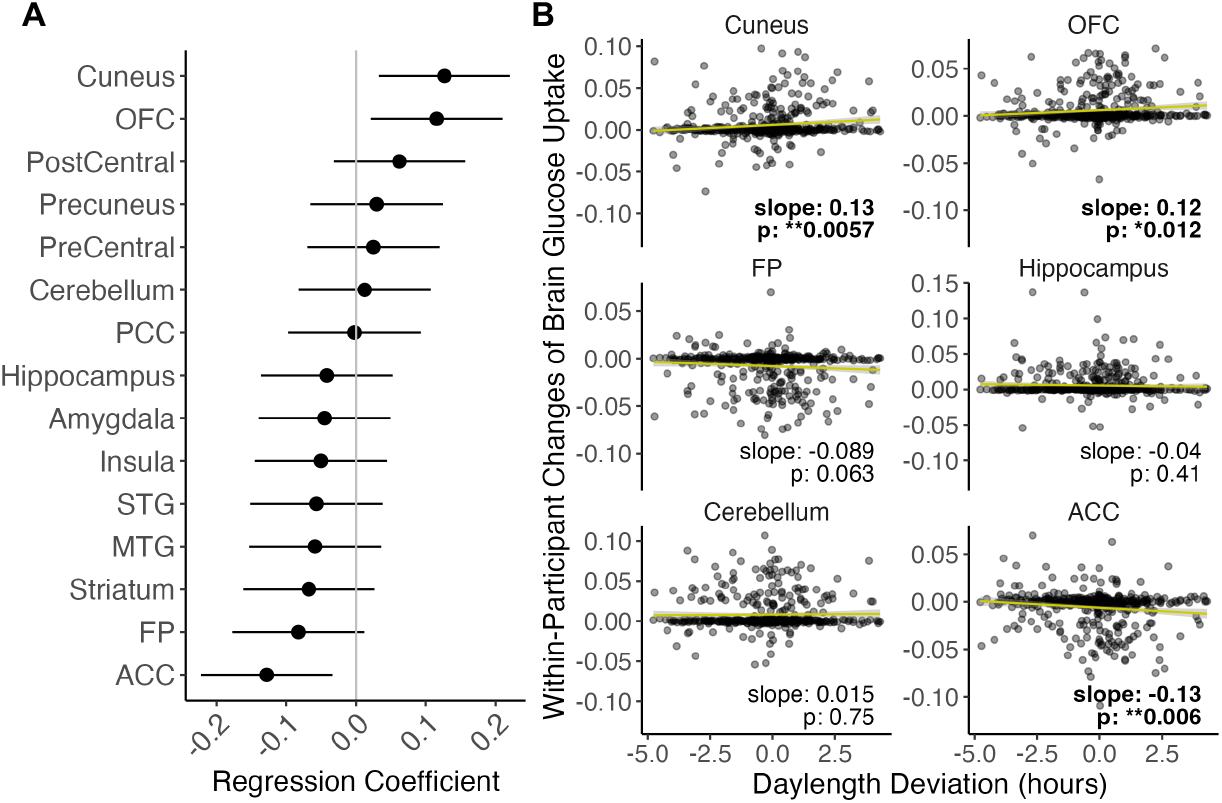
Repeated-measures analysis showing that within-participant changes in brain regional glucose uptake are affected by day length deviations. **A)** 95% confidence intervals for individual ROIs where within-participant changes in glucose uptake are affected by day length deviations. Standardized regression coefficients are plotted. **B)** Dot plots of the selected ROIs showing the impact of day length deviation on regional glucose uptake. Yellow line shows the least-squares (LS) regression line (y ~ day length deviation + BMI deviation) predicting glucose uptake changes, and the shaded area shows 95% confidence intervals. * = p <0.05,** = p <0.01. BMI, body mass index; ROI, region of interest.

Along with brain PET scans, day length was found to modulate multiple physiological features related to glucose uptake and immune responses. A longer day length was associated with reduced blood glucose and platelets levels in both the baseline and follow-up studies (**Fig. 5A**). Similarly, within-participant changes showed that an increase in day length was associated with reduced blood glucose levels and RBC and platelet counts (**Fig. 5B**).

**Figure 5.**
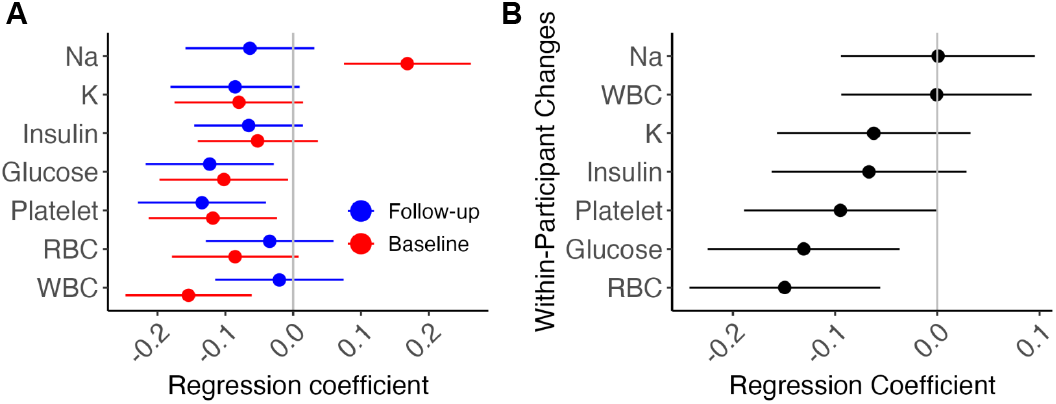
Effect of day length on physiological measures. **A)** Impact of day length on blood measurements at baseline and follow-up. **B)** Repeated-measures analysis showing the impact of within-participant daylength deviations on changes in physiological features. 95% confidence intervals are plotted, and standardized regression coefficients are used. WBC, white blood cell; RBC, red blood cell.

In addition, within-participant changes in BMI modulated regional glucose uptake and blood test results (**Supplementary Figure S4**). An increase in BMI was associated with reduced glucose uptake in the hippocampus, striatum, and amygdala. In the blood tests, a higher BMI was associated with increased insulin levels and WBC and RBC counts.

Day length had no statistically significant impact on stress or depression scores in either separate or within-participant data analysis.

## Discussion

The current study, based on repeated [^18^F]FDG-PET scans of a large cohort, provides a landmark evidence for photoperiodicity in glucose metabolism in the human brain. Our data analysis highlights that glucose consumption, especially in the cuneus, is consistently influenced by the photoperiod. Therefore, this study underscores the significance of photoperiodicity in human brain physiology, and the cuneus, anatomically adjacent to the primary visual cortex, may play a role in translating photoperiod information into temporal cues that regulate seasonal rhythms in the brain and its function.

In brain energy homeostasis, photoperiodicity is crucial for survival and well-being, as exemplified in extreme cases such as hibernation (22). In non-hibernating primates, the concept is often debated despite numerous reports on seasonal adaptations in gene transcription (23), neurotransmitter signaling (4, 5), immune response (24), and metabolic processes (6). Cross-sectional studies often face the drawback of uncontrolled individual variability, such as differences in socioeconomic status, although large sample sizes may help mitigate these issues and provide more reliable insights. The current study, which utilized longitudinal, repeated PET measurements from a large sample, provides significant evidence for the photoperiodicity of brain function, specifically concerning glucose consumption.

The cuneus was highlighted for its sensitivity to day length in the current study. It is located adjacent to the light-sensitive visual cortical regions that directly receive signals from the retina and is one of the areas immediately activated by visual stimuli. The anteromedial cuneus is thought to correspond to the homolog of the monkey area V6 (25). The cuneus is also a major component of the inferior fronto-occipital fasciculus, which originates in the cuneus, traverses the corpus callosum, and extends to the frontal lobes, including the superior frontal gyrus and orbital frontal cortex (26). This finding aligns with that of a previous cross-sectional study showing that a longer photoperiod is associated with higher glucose metabolic rates in the cuneus (6). Therefore, the cuneus may play a key role in translating photoperiodic changes in the environment into temporal cues that influence cognitive function and behavior.

In addition to the cuneus, the orbitofrontal cortex has also been identified to be modulated by photoperiod, as demonstrated by both separate ROI and repeated-measures analyses. However, this effect was largely absent in the full-volume analyses. The cuneus and orbitofrontal cortex are both key nodes in the social brain network, and their normal functions are closely linked to mental health (27), as well as social and feeding behaviors (28). Additionally, longer day length was associated with reduced glucose uptake in the anterior cingulate cortex. Full-volume analyses revealed that longer day length correlates with increased glucose consumption in the somatosensory area, superior parietal lobe, and mid-frontal and mid-temporal gyri—crucial regions for processing social and emotional cues (29). Moreover, a significant decrease in cerebellar glucose uptake was observed in both baseline and follow-up scans during longer photoperiods, although this finding was not replicated in the repeated-measures analysis. Overall, these results highlight the significant role of photoperiod in modulating glucose metabolism in brain regions that are vital for social cognition and behavior.

In addition to glucose metabolism in the brain, increased day length is associated with lower blood glucose levels, consistent with a previous report (30). Furthermore, the relationship between day length and platelet levels may reflect heightened inflammatory responses during shorter days. Shorter day length is hypothesized to drive energy expenditure to support the immune system during colder and darker periods (24). These observations further suggest that the human body is a finely tuned system, with coordinated brain and bodily responses to photoperiodic changes in the environment.

In the statistical analysis of the current study, BMI was used as a factor to control for the effects of the highly variable weight gain between scans, which is common during the dark season, on brain physiology. The data showed that within-participant changes in BMI are crucial modulators of subcortical limbic glucose metabolism. Increased BMI is associated with declined glucose consumption in the striatum and amygdala, underscoring the close relationship between the function of the brain reward circuitry and eating habits. This finding aligns with our previous research demonstrating that striatal glucose consumption directly influences food-reward responses of the brain and cognitive control (12). In addition, BMI changes within each individual affect blood insulin levels, whereas day length tends to modulate glucose levels. This suggests distinct mechanisms or a possible temporal mismatch between the two factors.

## Limitations

The database was compiled from historical scans in which relevant behavioral and self-reported measures, such as mood and eating habits, were poorly collected, resulting in significant missing data points. This study focuses on the potential photoperiodicity of human brain functions, deliberately excluding other environmental factors (e.g., atmospheric pressure and temperature) and social features (e.g., socioeconomic status and outdoor activities). In addition, daylight has multifaceted properties, including brightness, frequency, photoperiod, and day-to-day variations of the photoperiod, all of which may have detectable effects on brain function (1, 6). The current study focused on the role of photoperiod in directing brain function, as previously described (4, 5). Static PET scans, lacking efficient reference regions with minimal seasonal variation in glucose uptake, assume that global glucose uptake is least affected by photoperiod changes. However, this reference model may exaggerate photoperiod-related changes in some regions while producing reversed effects in others during group-level analysis. Nonetheless, within-participant comparisons effectively mitigate this limitation. Furthermore, the absence of structural brain MRI images limits our ability to distinguish between functional divisions in regional glucose signaling, such as gray and white matter signals.

## Conclusion

This study highlights the significant evidence of photoperiodicity in glucose metabolism in the human brain. We demonstrated the cuneus as a center for decoding photoperiod information; however, this finding requires further validation. In contemporary society, human circadian rhythms face continuous challenges owing to excessive exposure to artificial light, prolonged screen use, and irregular work shifts. The impact of these irregularities in light exposure on brain function, emotions, and social behavior may be significantly underestimated. A deeper understanding of inherent rhythms the brain could translate into knowledge that promotes healthier lifestyles and enhances well-being.

## Acknowledgement

This study was supported by the National Research Foundation of Korea (2020R1F1A1054201; KP), the National Natural Science Foundation of China (82272042; XL), and Fudan University affiliated with Huashan Hospital start-up funding (LS). XL received funding from the Pudong New Area Clinical Characteristic Discipline Project (No. PWYts2021-01), Clinical Research Program of Health Industry of Shanghai Municipal Commission of Health (202150002), Pudong Hospital, Fudan University, College Level Project (YJYJRC202108/ YJYJRC202101/Zdzk2020-14).

## Data Availability

Physiological and behavioral data of the human brain are available upon request from KP.

## Author Contributions

KP and LS conceptualized the study and conducted data analysis. All authors contributed to the interpretation of the data and writing of the manuscript.

