## Supplementary for "Photoperiodicity in Glucose Metabolism in the Human Brain"

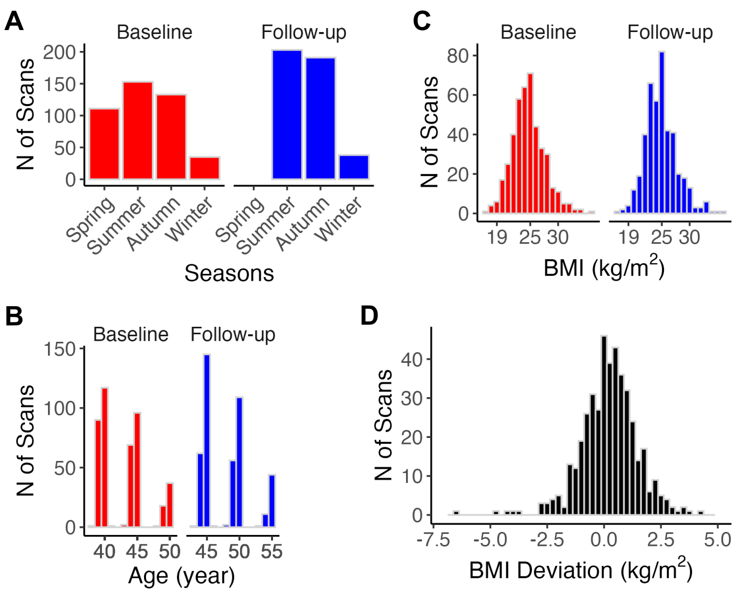


***Supplementary Figure S1.*** *Distribution of data variables. Distribution of seasons* ***(A)****, BMI* ***(B)*** *and age* ***(C)*** *in the baseline and follow-up data. One bin represents 1 kg/m$^2$ BMI or 1 year of age.* ***(D)*** *Distribution of with-participant deviation in BMI between repeated scans. One bin represents a BMI of 0.25 kg/m$^2$. BMI, body mass index.*

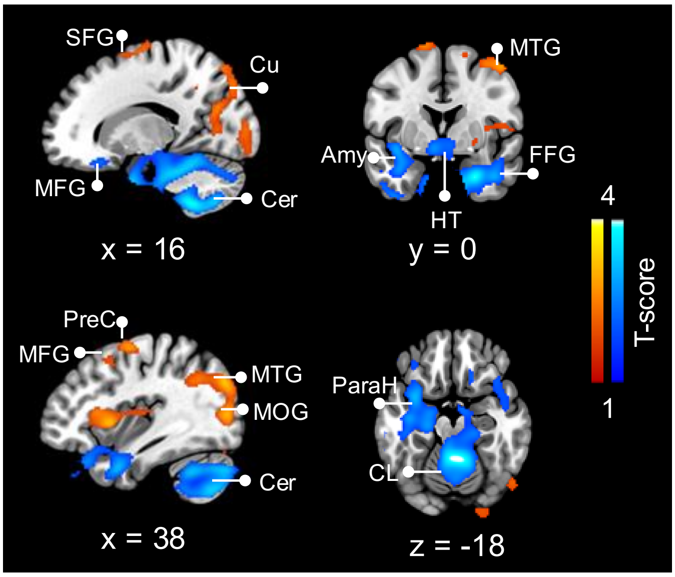


***Supplementary Figure S2.*** *Brain regions where glucose uptake is sensitive to day length in follow-up scans. Baseline measures demonstrate positive (hot color) and negative (cold color) linear relationships with day length. Data are FDR cluster-level corrected at p < 0.05. Amy, amygdala; Cer, cerebellum; CL, cerebellar lingual gyrus; Cu, cuneus; FFG, fusiform gyrus; HT, hypothalamus; MFG, middle frontal gyrus; MOG, middle occipital gyrus; MTG, middle temporal gyrus; ParaH, parahippocampus; PreC, precentral gyrus; SFG, superior frontal gyrus.*

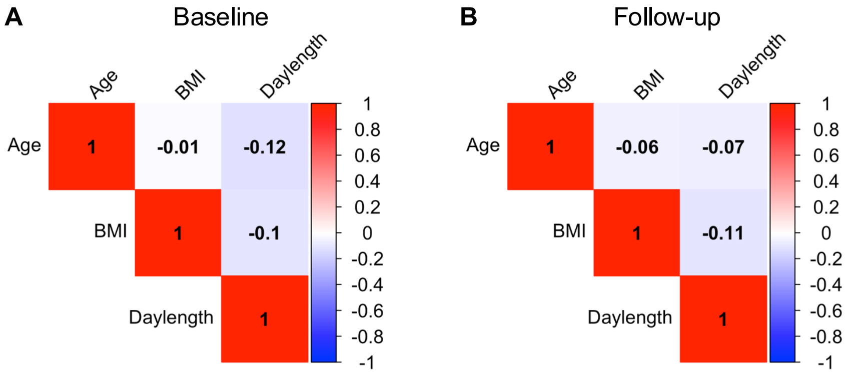


***Supplementary Figure S3.*** *Correlation coefficients between age, BMI, and day length in* ***(A)*** *baseline and* ***(B)*** *follow-up scans across participants. BMI, body mass index.*

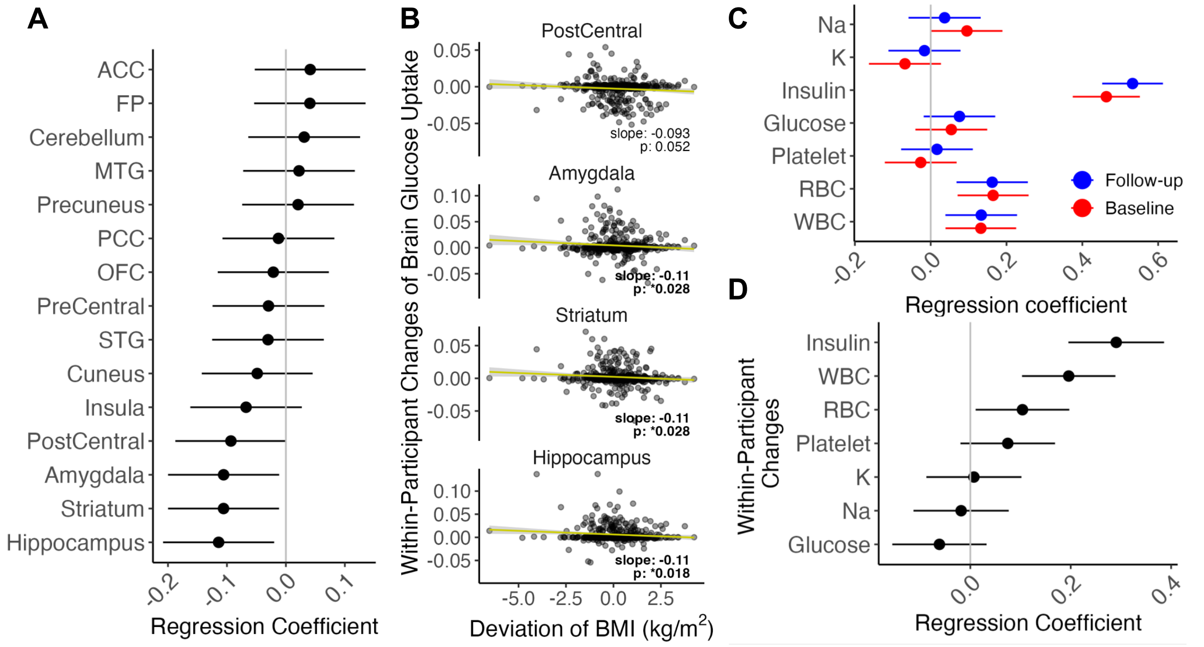


***Supplementary Figure S4.*** *Effect of with-participant BMI deviation on brain glucose uptake and blood measurements.* ***(A)*** *ROIs where with-participant changes in glucose uptake are affected by with-participant deviation in BMI.* ***(B)*** *Dot plots of selected ROIs showing the impact of with-participant BMI deviation.* ***(C)*** *Impact of BMI deviation on blood measurements, as indicated by both separate data analysis and* ***(D)*** *with-participant comparisons. 95% confidence intervals are plotted, and standardized regression coefficients are used. BMI, body mass index; ROI, region of interest.*

***Supplementary Table S1.*** *Participant characteristics of the baseline and follow-up studies.*

| | Baseline study | Follow-up study | p |
|---|---|---|---|
| Age (years) | 42.8±3.6 | 47.9±3.5 | <0.0001 |
| BMI (kg/m$^2$) | 24.7±2.9 | 24.9±2.8 | <0.0001 |
| Blood test | | | |
| White blood cell (x $10^9$/L) | 6.4±1.6 | 6.1±1.5 | <0.0001 |
| Red blood cell (x $10^{12}$/L) | 5.0±0.3 | 5.1±0.4 | 0.0012 |
| Platelet (x $10^9$/L) | 246.0±52.3 | 262.5±63.5 | <0.0001 |
| Glucose (mg/dL) | 93.0±16.6 | 101.9±24.1 | <0.0001 |
| Insulin (uIU/mL) | 5.3±3.1 | 6.8±3.9 | <0.0001 |
| Na (mmol/L) | 142.5±1.8 | 142.2±1.8 | 0.0592 |
| K (mmol/L) | 4.5±0.4 | 4.5±0.3 | 0.4393 |
| Marital status | | | 0.2844 |
| Married | 389 | 404 | |
| Single | 14 | 12 | |
| Divorced | 5 | 12 | |
| Widowed | 0 | 1 | |
| Education level (years) | 13.5±2.1 | 13.6±2.1 | 0.0031 |
| Questionnaires | | | |
| Depression | 4.9±4.7 | 5.1±3.8 | 0.1364 |
| Stress | 16.0±6.2 | 15.1±6.1 | 0.0021 |

Mean ± standard deviation; BMI, body mass index.

***Supplementary Table S2****. Comparison of participant characteristics in groups with longer or shorter day length scans.*

| | Longer daylength | Shorter daylength | p |
|---|---|---|---|
| BMI (kg/m$^2$) | 24.7±2.9 | 24.8±2.8 | 0.2001 |
| Day length (hours) | 14.1±1.4 | 12.8±1.5 | <0.0001 |
| No. of scans from baseline study | 236 | 196 | |
| No. of scans from follow-up study | 196 | 236 | |
| Marital status | | | |
| Married | 396 | 397 | |
| Single | 15 | 11 | |
| Divorced | 8 | 9 | |
| Widowed | 0 | 1 | |
| Education level (years) | 13.6±2.1 | 13.6±2.0 | 0.3086 |
| Questionnaires | | | |
| Depression | 4.7±4.0 | 5.3±4.5 | 0.9797 |
| Stress | 15.6±6.0 | 15.5±6.3 | 0.3595 |

BMI, body mass index.
